# Clinical Validation of a Non-Invasive Prenatal Test for Genome-Wide Detection of Fetal Copy Number Variants

**DOI:** 10.1101/033555

**Authors:** Roy B. Lefkowitz, John A. Tynan, Tong Liu, Yijin Wu, Amin R. Mazloom, Eyad Almasri, Grant Hogg, Vach Angkachatchai, Chen Zhao, Daniel S. Grosu, Graham McLennan, Mathias Ehrich

## Abstract

1.

**Background:** Current cell-free DNA (cfDNA) assessment of fetal chromosomes does not analyze and report on all chromosomes. Hence, a significant proportion of fetal chromosomal abnormalities are not detectable by current non-invasive methods. Here we report the clinical validation of a novel NIPT designed to detect genome-wide gains and losses of chromosomal material ≥7 Mb and losses associated with specific deletions <7 Mb.

**Objective:** The objective of this study is to provide a clinical validation of the sensitivity and specificity of a novel NIPT for detection of genome-wide abnormalities.

**Study Design:** This retrospective, blinded study included maternal plasma collected from 1222 study subjects with pregnancies at increased risk for fetal chromosomal abnormalities that were assessed for trisomy 21 (T21), trisomy 18 (T18), trisomy 13 (T13), sex chromosome aneuploidies (SCAs), fetal sex, genome-wide copy number variants (CNVs) 7 Mb and larger, and select deletions smaller than 7 Mb. Performance was assessed by comparing test results with findings from G-band karyotyping, microarray data, or high coverage sequencing.

**Results:** Clinical sensitivity within this study was determined to be 100% for T21, T18, T13, and SCAs, and 97.7% for genome-wide CNVs. Clinical specificity within this study was determined to be 100% for T21, T18, and T13, and 99.9% for SCAs and CNVs. Fetal sex classification had an accuracy of 99.6%.

**Conclusion:** This study has demonstrated that genome-wide non-invasive prenatal testing (NIPT) for fetal chromosomal abnormalities can provide high resolution, sensitive, and specific detection of a wide range of sub-chromosomal and whole chromosomal abnormalities that were previously only detectable by invasive karyotype analysis. In some instances, this NIPT also provided additional clarification about the origin of genetic material that had not been identified by invasive karyotype analysis.

## 2. INTRODUCTION

Since its introduction in 2011, NIPT has had a significant impact on prenatal care. In only four years, NIPT has evolved into a standard option for high-risk pregnancies [1]. Content has also evolved from exclusive T21 testing to include T18, T13, SCAs, and select microdeletions. This ‘standard’ content can be expected to detect 80–83% of chromosomal abnormalities detected by karyotyping in a general screening population [2–4], however this leaves a gap of approximately 17–20% of alternative chromosomal/sub-chromosomal abnormalities not detected. Consequently, obtaining comprehensive information about the genetic makeup of the fetus requires an invasive procedure. To overcome these limitations, NIPT should be extended to cover the entire genome. However, it is challenging to maintain a very high specificity and positive predictive value when interrogating all accessible regions in the genome [5]. In previous reports, we have overcome these technical hurdles [6]. Furthermore, a recent study by Yin et al. demonstrated feasibility for non-invasive genome-wide detection of sub-chromosomal abnormalities [7]. In this report, we have improved the assay and the statistical methods to enable comprehensive genome-wide detection of CNVs ≥7 Mb. We present the results of a large blinded clinical study of more than 1200 samples including more than 100 samples with common aneuploidies detectable by traditional NIPT and over 30 samples affected by sub-chromosomal CNVs.

## 3. METHODS

### STUDY DESIGN

This blinded, retrospective clinical study included samples from women considered at increased risk for fetal aneuploidy based on advanced maternal age ≥35, a positive serum screen, an abnormal ultrasound finding, and/or a history of aneuploidy. Archived samples were selected for inclusion in the study by an unblinded internal third party according to the requirements documented in the study plan. The samples were then blind-coded to all operators and the analysts who processed the samples. After sequencing, an automated bioinformatics analysis was performed to detect whole chromosome aneuploidies and sub-chromosomal CNVs. The results were compiled electronically and were reviewed by a subject matter expert who assigned the final classification. This manual review mimics the process in the clinical laboratory, where cases are reviewed by a laboratory director before a result is signed out. The complete list of classification results was provided to the internal third party for determination of concordance. Analyzed samples had confirmation of positive or negative events by either G-band karyotype or microarray findings from samples collected through either CVS or amniocentesis. Circulating cell-free "fetal” DNA is believed to originate largely from placental trophoblasts. Genetic differences between the fetus and the placenta can occur (e.g. confined placental mosaicism), leading to discordance between NIPT results and cytogenetic studies on amniocytes or postnatally obtained samples [8]. Results from CVS by chromosomal microarray were thus considered the most accurate ground truth. Therefore, discordant results originating from amniocytes (karyotype or microarray) were resolved by sequencing at high coverage (an average of 226 million reads per sample). Sequencing depth has been shown as the limiting factor in NIPT methods, with increased depth allowing improved detection of events in samples with lower fetal fractions or improved detection of smaller events [9]. High coverage sequencing has been used in multiple studies to unambiguously identify sub-chromosomal events ([6,10–12]) and was used here as a reference for performance evaluation in discrepant amniocentesis samples.

Details of the sample demographics are described in Table 1. Indications for invasive testing are described in Table 2.

**Table 1.** Demographic and pregnancy-related data. Choice of invasive procedure, choice of diagnostic test, and demographic data were not available for all 1208 samples included in the study. Ntotal refers to the number of samples where that data was available.

| Demographic | Median | Range |
| --- | --- | --- |
| Maternal Age (Yrs) (Ntotal = 1177) | 36.0 | 17.8-47 |
| Gestational Age (Wks) (Ntotal =1183) | 17 | 8-38 |
| Maternal Weight (Lbs) (Ntotal =1168) | 150 | 93-366 |
| Procedure | Percent | (Naffected/Ntotal) |
| CVS | 14.5 | (175/1203) |
| Amniocentesis | 85.2 | (1025/1203) |
| Both | 0.2 | (3/1203) |
| Confirmation | Percent | (Naffected/Ntotal) |
| Karyotype | 90.4 | (1089/1205) |
| Microarray | 5.8 | (70/1205) |
| Both | 3.8 | (46/1205) |

**Table 2.** Indication for testing. Each cell lists the count and percentage of different indications for testing for different types of chromosomal abnormalities. Samples may have multiple indications for testing. This table includes both reportable (n=1166) and non-reportable samples (total n=42), for a total of 1208 samples. One sample had no indication for testing, leaving a total of 1207 samples for the above analysis. As 2 of the trisomy 18 samples also had an SCA, the total number of abnormalities and euploid samples sum up to 1209.

| Indication for Testing | Euploid<br>(n=1009) | T21<br>(n=87) | T18<br>(n=29) | T13<br>(n=15) | SCA<br>(n=26) | CNV<br>(n=43) | Total<br>(n=1209) |
| --- | --- | --- | --- | --- | --- | --- | --- |
| Positive Serum Screening | 404(40%) | 36(41.4%) | 13(44.8%) | 4(26.7%) | 3(11.5%) | 16(37.2%) | 476(39.4%) |
| Maternal Age > 35 Yrs | 418(41.4%) | 28(32.2%) | 10(34.5%) | 4(26.7%) | 2(7.7%) | 10(23.3%) | 472(39%) |
| Ultrasound Abnormality | 152(15.1%) | 34(39.1%) | 6(20.7%) | 10(66.7%) | 16(61.5%) | 21(48.8%) | 239(19.8%) |
| Positive Family History | 33(3.3%) | 4(4.6%) | 1(3.4%) | 0(0%) | 1(3.8%) | 1(2.3%) | 40(3.3%) |
| Not Specified | 85(8.4%) | 11(12.6%) | 5(17.2%) | 2(13.3%) | 4(15.4%) | 2(4.7%) | 109(9%) |

### SAMPLE COLLECTION

In total, 1222 maternal plasma samples had been previously collected using Investigational Review Board (IRB) approved protocols (protocol numbers: Compass IRB #00508, WIRB #20120148, Compass IRB #00351, Columbia University IRB #AAAN9002) with a small subset (9 samples) comprising remnant plasma samples collected from previously consented patients in accordance with the FDA Guidance on Informed Consent for *In Vitro* Diagnostic Devices Using Leftover Human Specimens that are Not Individually Identifiable (25 April 2006). All subjects provided written informed consent prior to undergoing any study related procedures.

### LIBRARY PREPARATION, SEQUENCING, AND ANALYTICAL METHODS

Libraries were prepared and quantified as described by Tynan et al.[13]. To reduce noise and increase signal, sequencing depth for this analysis was increased to target 32M reads per sample. Sequencing reads were aligned to hg19 using Bowtie 2 [14]. The genome was then partitioned into 50-kbp non-overlapping segments and the total number of reads per segment was determined, by counting the number of reads with 5’ ends overlapping with a segment. Segments with high read-count variability or low map-ability were excluded. The 50 kbp read-counts were then normalized to remove coverage and GC biases, and other higher-order artifacts using the methods previously described in Zhao et al [6].

The presence of fetal DNA was quantified using the regional counts of whole genome single-end sequencing data as described by Kim et al. [15].

### GENOME-WIDE DETECTION OF ABNORMALITIES

Circular binary segmentation (CBS) was used to identify CNVs throughout the entire genome by segmenting each chromosome into contiguous regions of equal copy number [16]. A segment-merging algorithm was then used to compensate for over segmentation by CBS when the signal-to-noise ratio (SNR) was low [17]. Z-scores were calculated for both CBS-identified CNVs and whole chromosome variants by comparing the signal amplitude with a reference set of samples in the same region. The measured Z-scores form part of an enhanced version of <u>C</u>hromosomal <u>A</u>berration <u>DE</u>cision <u>T</u>ree (CADET) previously described in detail in Zhao et al [6]. CADET incorporates the Z statistics for a CBS-detected CNV to assess the its statistical significance and a log odds ratio (LOR) to provide a measure of the likelihood of an event being real, based on an observed fraction of fetal DNA across the genome (see supplemental methods for [6]).

To further improve specificity of CNV detection, bootstrap analysis was performed as an additional measure for the confidence of the candidate CNVs. The within sample read count variability was compared to a normal population (represented by 371 euploid samples) and quantified by bootstrap confidence level (BCL). In order to assess within sample variability, the bootstrap resampling described below was applied to every candidate CNV.

For each identified segment within the CNV, the median shift of segment fraction from the normal level across the chromosome was calculated. This median shift was then corrected to create a read count baseline for bootstrapping. Next, a bootstrapped segment of the same segment length as the candidate CNV was randomly sampled with replacement from the baseline read counts. The median shift was then applied to this bootstrapped fragment. The segment fraction of the bootstrapped fragment was calculated as follows:

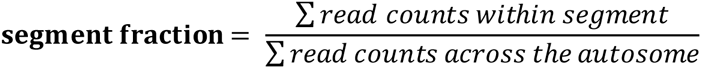

This process was repeated 1000 times to generate a bootstrap distribution of segment fractions for an affected population. A normal reference distribution was created based on the segment fraction of the same location as the candidate CNV in 371 euploid samples. A threshold was then calculated as the segment fraction that was at least 3.95 median absolute deviations away from the median segment fraction of the reference distribution. Lastly, the BCL was calculated as the proportion of bootstrap segments whose fractions had absolute z-statistics above the significance threshold.

A whole chromosome or sub-chromosomal abnormality is detected as follows:

1. A chromosome is classified to have a trisomy or monosomy if:

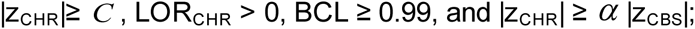
2. A chromosome is classified to have a sub-chromosomal abnormality if:

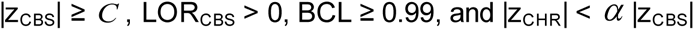
where *C* is a predefined z-score cutoff that controls the tradeoff between sensitivity and specificity. For this study *C* was 3 for chromosome 21 trisomy and monosomy, and 3.95 for all other CNVs. The comparison |z_CHR_| ≥ *α* |z_CBS_| is used to distinguish a whole chromosome event from a sub-chromosomal event. *α* denotes the type 1 error for misclassification of abnormalities as an aneuploidy. Simulations (see supplemental methods for [6]) showed that *α* = 0.8 resulted in a misclassification of abnormalities at close to 0%; as such, this value was used for this study.

## 4. RESULTS

The study comprised a total of 1222 maternal plasma samples. After unblinding, eleven samples were excluded because they had no or insufficient karyotype or microarray information (see Figure 1) and three samples were excluded because of confirmed mosaicism.

**Figure 1.**
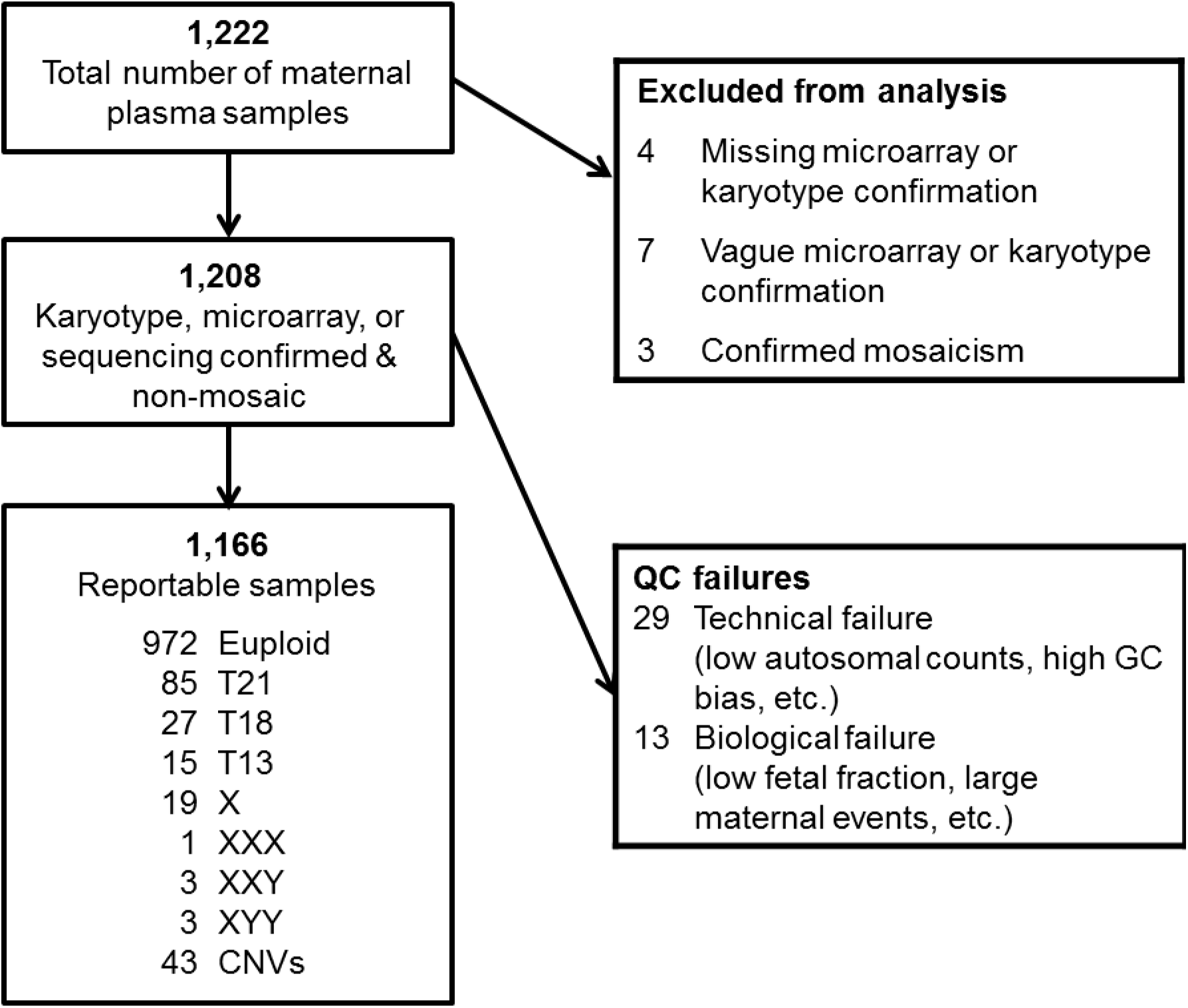
Flow chart showing sample exclusions and reasons for non-reportable samples. One of the trisomy 18 samples was also an XXY sample and one of the other trisomy 18 samples was also an X sample. For samples that had discordant results for sequencing versus karyotype or microarray derived from testing of amniocytes, uniplex sequencing was used for confirmation.

Forty-two (42) of the remaining 1208 samples were flagged as non-reportable using quality criteria that had been established prior to analysis, leaving 1166 reportable samples for analyses. Technical failure criteria included but were not limited to: low library concentration, low raw autosomal counts, high GC bias, poor normalization, and high bin variability (see Figure 1). Biological failure criteria included low fetal fraction (less than 4%) and large maternal CNV events. The most common reason for failure was low fetal fraction (n= 11). During review of the data, one sample was signed out as T18 (and was included in the analyzed cohort), even though it did not meet the autosomal count minimum. This sample had sufficient counts for the determination of standard aneuploidies, but not sufficient counts for the detection of sub-chromosomal CNVs. The 42 non-reportable samples showed no evidence of enrichment for whole-chromosome/sub-chromosomal positive samples (5 positive of 42 non-reportable samples vs. 204 positive of 1166 reportable samples). Concordant with previous studies, the overall reportable rate on first aliquots of maternal plasma was 96.5%.The overall no call rate per patient is expected to be approximately 1% when a second aliquot is available, based on published clinical laboratory experience [18].

### T21, T18, and T13 Detection

Among the 1166 reportable samples, there were 85 T21 samples, 27 T18 samples, and 15 T13 samples (see Figure 1). All euploid, T21, T18, and T13 samples (determined by invasive diagnostic procedures) were classified correctly by NIPT. One sample was classified by NIPT as T21 but had a normal (46, XX) karyotype by amniocentesis (see Supplemental Table 1). This discrepancy was adjudicated through high coverage ‘uniplex’ sequencing (typically with >180 million reads) according to our study plan (see Supplemental Figure 1D). The sample showed 16.3% fetal fraction, but the gain of genetic material from chromosome 21 was concordant with 6.5% fetal fraction. This is suggestive of confined placental mosaicism, a relatively common cause of discordant results between amniocyte results and CVS or placentally-derived cell free DNA analysis [8]. Table 3 summarizes the performance for T21, T18, and T13. For T21, the sensitivity was 100% (95% CI: 94.6%-100%) and the specificity was 100% (95% CI: 99.6%-100%). For T18, the sensitivity was 100% (95% CI: 84.4%-100%) and the specificity was 100% (95% CI: 99.6%-100%). For T13, the sensitivity was 100% (95% CI: 74.7%-100%) and the specificity was 100% (95% CI: 99.6%-100%).

**Table 3.** Clinical performance. Concordance was established using confirmation from a combination of microarray, G-band karyotype, and high coverage sequencing data. There were 42 samples that were non-reportable for all chromosomal abnormalities based on the breakdown in Figure 1. However, this table also takes into account additional reportability criteria for SCA based on chromosome X and Y Z-scores.

| Abnormality | Concordant Positive | Discordant Positive | Concordant Negative | Discordant Negative | Sensitivity | Specificity |
| --- | --- | --- | --- | --- | --- | --- |
| T21 | 85 | 0 | 1081 | 0 | 100%<br>(95% CI: 94.6%-100%) | 100%<br>(95% CI: 99.6%-100%) |
| T18 | 27 | 0 | 1139 | 0 | 100%<br>(95% CI: 84.4%-100%) | 100%<br>(95% CI: 99.6%-100%) |
| T13 | 15 | 0 | 1151 | 0 | 100%<br>(95% CI: 74.7%-100%) | 100%<br>(95% CI: 99.6%-100%) |
| SCA | 26 | 1 | 1117 | 0 | 100%<br>(95% CI: 84.0%-100%) | 99.9%<br>(95% CI: 99.4%-100%) |
| CNVs* | 42 | 1 | 1122 | 1 | 97.7%<br>(95% CI: 86.2%-99.9%) | 99.9%<br>(95% CI: 99.4%-100%) |
| *- Genome copy number variants including 8 samples with detected whole chromosome trisomies, and 35 samples with sub-chromosomal CNVs |  |  |  |  |  |  |

**Table 3. Clinical performance.**
| Abnormality | Concordant Male | Discordant Male | Concordant Female | Discordant Female | Accuracy |
| --- | --- | --- | --- | --- | --- |
| Fetal Sex | 583 | 4 | 578 | 1 | 99.6%<br>(95% CI: 98.9%-99.8%) |

### SCA Detection

In the 1166 samples reportable for all chromosomal abnormalities, there were 21 samples that were flagged as non-reportable specifically for SCA classification based on thresholds for the chromosome Xz-score and chromosome Yz-score as described by Mazloom et al. [19]. There was also one additional sample with an apparent maternal XXX that was flagged as non-reportable for SCA because the maternal event distorted the sex chromosomal representation to a degree that fetal events could not be classified. Among the remaining 1144 samples reportable for SCAs, there were 7 discordant positives that were classified as normal (46, XX) by karyotype and as XO by sequencing at 6-plex (see Supplemental Table 1). In all discordant cases, the karyotype had been obtained from amniocyte samples. This phenomenon is well described and may be attributed to varying levels of placental or maternal mosaicism [20]. Uniplex sequencing confirmed 6 of the 7 XO samples. The 7^th^ sample had a non-reportable result at uniplex coverage, hence the existing amniocentesis result was used as truth resulting in one false positive assignment. Overall, the sensitivity for SCA was 100% (95% CI: 84.0%-100%) with a specificity of 99.9% (95% CI: 99.4%-100%) (see Table 3).

### Genome Wide Detection of CNVs

The test was also designed to detect CNVs equal to or larger than 7 Mb (including whole chromosome abnormalities other than T13, T18, T21, and SCAs); as well as select microdeletions smaller than 7 Mb [5]. Among the 1166 samples reportable for sub-chromosomal abnormalities, there were 43 samples that had positive results for a variety of CNV aberrations: Wolf-Hirschhorn syndrome deletions, DiGeorge syndrome deletions, Prader-Willi/Angelman syndrome deletions, Cri du Chat syndrome deletions, and a variety of ≥7 Mb CNVs including both sub-chromosomal CNVs and whole chromosome trisomies. Several examples of NIPT detected CNVs confirmed by microarray or karyotype are shown in Figure 2.

**Figure 2.**
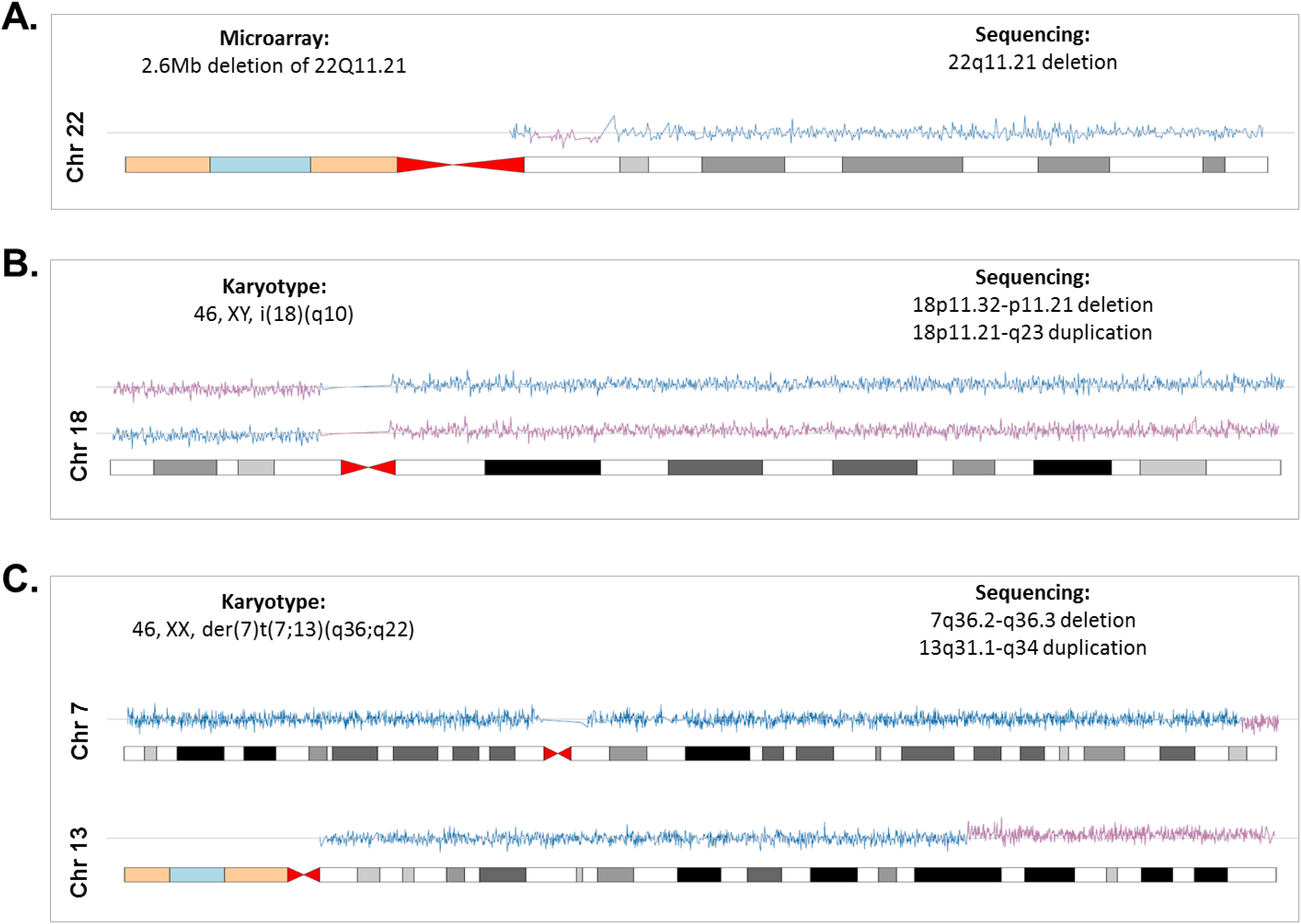
Examples of karyotype- and microarray-detected CNVs detected by sequencing. A) Chromosome 22 ideogram showing sequencing-based detection of a 22q11.21 deletion (DiGeorge syndrome) that was confirmed by karyotype and microarray. B) Chromosome 18 ideogram for a sample with a karyotype 46, XY, i(18)(q10). C) Chromosome 7 and 13 ideograms for a sample with a karyotype 46, XX, der(7)t(7;13)(q36;q22).

Overall, the sensitivity for detection of whole chromosome and sub-chromosomal abnormalities other than T13, T18, T21, and SCAs was 97.7% (95% CI: 86.2%-99.9%) and the specificity was 99.9% (95% CI: 99.4%-100%) (see Table 3). One case was clearly mosaic for T22 by both standard coverage and uniplex sequencing (see Supplemental Table 1 and Supplement Figure 1E), but was classified as normal by microarray analysis. Because the invasive diagnosis came from microarray analysis of cells derived from CVS, our study design considered this as the gold standard, and this outcome was considered a discordant positive. Another case showed no gain or loss of genetic material with both standard and uniplex sequencing for a sample that had a 46, XX, der(12)t(12;19)(p13.1;q13.1) karyotype (see Supplemental Figure 2B). This outcome was considered as a discordant negative given that the invasive procedure was CVS. A set of 7 samples were classified as full chromosomal trisomies, and because the karyotype or microarray results were derived from amniocytes in these cases, uniplex sequencing was performed for adjudication per the study design. The trisomy finding was confirmed in each case by uniplex sequencing, and these findings were considered as concordant positives (Supplemental Figure 1).

In addition to the high accuracy demonstrated in the results, in some cases NIPT could also provide clarification about the origin of extra genetic material when the G-banding pattern was not sufficiently clear. In one case, the amniocyte karyotype finding, 46, XX, der(5)t(5;?)(p15.3;?), indicated a deletion on chromosome 5 and a duplication of unknown origin (see Figure 3). NIPT identified an 18.6 Mb duplication representing the entire short arm of chromosome 17, indicating this fetal tissue possessed a trisomy of chromosome 17p that was not clarified by standard karyotype. Trisomy 17p is associated with developmental delay, growth retardation, hypotonia, digital abnormalities, congenital heart defects, and distinctive facial features [21]. Additional cases are shown in Supplemental Figures 3.

**Figure 3.**
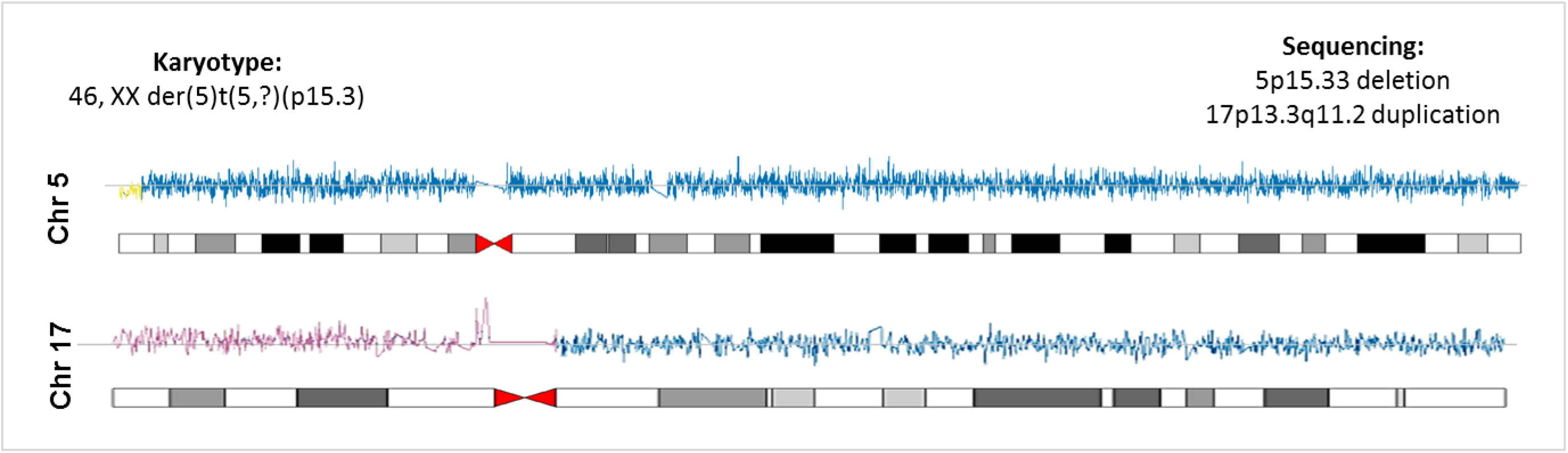
Clarification of karyotype findings by sequencing of maternal plasma. Normal regions are colored blue while deletions and duplications that are detected with high bootstrap confidence levels (>0.99) are colored red. Low bootstrap confidence events are colored yellow. Chromosome 5 & 17 ideograms are shown for a sample with a karyotype of 46, XX, der(5)t(5;?)(p15.3;?). The unidentified duplication on the karyotype is from chromosome 17.

## 5. DISCUSSION

Couples and families who are seeking prenatal information about the genetic health of their baby have multiple options today – ranging from non-invasive tests that screen for select chromosomal abnormalities to invasive procedures including karyotype and microarray testing that can deliver the most comprehensive genomic assessment. Current NIPT methods provide information about a limited set of conditions, that typically include T21, T18, T13 and SCAs. In this study we expand testing to the entire genome. The results demonstrate that high sensitivity and specificity were maintained while adding genome-wide clinically relevant content.

Most NIPT tests suffer from the problem of multiple hypothesis testing; when multiple regions of the genome are tested independently, false positive rates become additive [22]. This can result in unacceptably high false positive rates and lead to maternal anxiety and additional unnecessary clinical testing. Methods that target individual regions of the genome cannot overcome these limitations because they are part of the design of the test [23]. We have shown that these limitations can be overcome by using a genome-wide approach, as described here and previously [6]. In this study we demonstrate that the test can provide excellent performance for CNVs ≥7Mb and select microdeletions <7Mb, across the genome.

This study has two main limitations. Although we analyze 2- to 6-fold more samples with sub-chromosomal CNVs than previous studies [6, 11, 23], the total number of affected samples is still relatively small compared to studies validating performance for detection common whole-chromosome aneuploidies [24]. It remains challenging to statistically validate detection of rare chromosomal variations. A second challenge commonly observed in novel technologies with increased performance is the compilation of samples with appropriate ground truth. In our study we face the technical challenge of comparing sequencing versus G-band karyotype or microarray, as well as the biological aspect of cfDNA versus amniocyte or CVS. We have used a method that prioritizes CVS over amniocentesis because of the placental origin of cell free DNA [8]. Factors that can lead to discordant results include confined placental mosaicism, maternal mosaicism, vanishing twins, undetected tumors, and technical errors.

An interesting observation in this study was the sometimes subjective nature of karyotyping. When G-banding is used as an analytical method, the optical resolution is occasionally not sufficient to determine the origin of additional genetic material, making a clinical interpretation more difficult. These ambiguities are eliminated when using NIPT because the test requires sequencing alignment to the genome before DNA gains and losses are determined. However, G-banding will, for the foreseeable future, be the superior methodology to determine copy number neutral structural changes.

In pregnancies that can benefit from additional information, this test provides more clinically relevant results than previous NIPT options. However, its role as a follow-up test to abnormal ultrasound findings or as a general population screen will likely be debated for the foreseeable future. It has been argued that the incidence of clinically relevant sub-chromosomal CNVs could be as high as 1.7% in a general population [25], and early microarray data [26] indicate that as many as 30% of sub-chromosomal CNVs could be >7Mb. In consequence, the incidence for these large CNVs may be around 0.5% in the general population, substantially higher than the incidence of T21.

In summary, this clinical study provides validation for an approach that extends the clinical validity of cfDNA testing, now providing detection of T21, T18, T13, SCAs, fetal sex, and genome-wide detection of sub-chromosomal and whole chromosomal abnormalities. Overall, sub-chromosomal abnormalities and aneuploidies other than T13, T18, T21, and SCAs were detected with a combined clinical sensitivity and specificity of 97.7% and 99.9%, respectively. This enables comprehensive non-invasive chromosomal assessment that was previously available only by karyotype, and in some cases, may clarify cryptic findings otherwise identifiable only by microarray.

## Acknowledgements

The authors would like to acknowledge the efforts of those who aided completion of this study: Nathan Faulkner and Ashley Diaz for sample procurement; Sarah Ahn, Christine Safanayong, Haiping Lu, Joshua White, Lisa Chamberlain for sample processing and DNA extraction; David Wong, Jana Schroth, Marc Wycoco, Jesse Fox for library preparation and sequencing, Pat Liu for bioinformatics pipeline processing. We would also like to acknowledge the contribution of samples by Ronald Wapner at Columbia University.

